# BERMUDA: A novel deep transfer learning method for single-cell RNA sequencing batch correction reveals hidden high-resolution cellular subtypes

**DOI:** 10.1101/641191

**Authors:** Tongxin Wang, Travis S Johnson, Wei Shao, Zixiao Lu, Bryan R Helm, Jie Zhang, Kun Huang

## Abstract

To fully utilize the power of single-cell RNA sequencing (scRNA-seq) technologies for cell lineation and identifying *bona fide* transcriptional signals, it is necessary to combine data from multiple experiments. We present *BERMUDA* (Batch-Effect ReMoval Using Deep Autoencoders) — a novel transfer-learning-based method for batch-effect correction in scRNA-seq data. *BERMUDA* effectively combines different batches of scRNA-seq data with vastly different cell population compositions and amplifies biological signals by transferring information among batches. We demonstrate that *BERMUDA* outperforms existing methods for removing batch effects and distinguishing cell types in multiple simulated and real scRNA-seq datasets.

## Background

Single-cell transcriptional dynamics are important for understanding molecular physiology and disease dysregulation within heterogeneous tissues. Until recently, the standard techniques for single-cell analysis were flow cytometry [1, 2] and fluorescence imaging of tissue on slides [3, 4]. Though these techniques have provided tremendous insights, they are limited to a small, pre-defined set of molecular markers [1]. More recently, high-throughput techniques such as RNA sequencing (RNA-seq) were established to measure expression of thousands of genes, but were designed for bulk tissue samples [5, 6]. Single-cell RNA sequencing (scRNA-seq) was developed to characterize high-throughput gene expression profiles for populations of individual cells, which has enabled an unprecedented resolution of cellular heterogeneity in complex tissues. Widespread adoption of scRNA-seq techniques have produced large complex datasets, which present new computational challenges for evaluating experimental reproducibility and combining data from different batches and platforms [5, 7–11].

There have been many attempts to combine gene expression data from different experiments to achieve a more comprehensive understanding of the underlying cellular heterogeneity. The first generation of tools were adapted from linear model analysis of microarrays [12–14] and were subsequently modified for RNA-seq data via generalized linear [15] or negative binomial models [16]. These methods represent the foundation of batch-effect removal when scRNA-seq data from different sequencing runs are combined; however, scRNA-seq poses additional challenges when combining disparate data. Unlike microarray or whole-tissue RNA-seq, scRNA-seq is especially prone to “drop out” events in which RNA is not amplified during library preparation [17, 18]. Cell types and proportions may vary substantially across samples [19, 20]. Both technical and biological variability contribute to strong batch effects (*i.e*., systematic variance) that must be overcome to meaningfully combine datasets as is fundamental in comparative and bioinformatic studies [21, 22].

*Seurat-CCA* (*Seurat v2*) [22] and *mnnCorrect* [20] were the first methods proposed to combine scRNA-seq data from multiple batches. *Seurat v2* uses canonical correlation analysis (CCA) to project cells from different experiments to a common bias-reduced low-dimensional representation. However, this type of correction does not account for the variations in cellular heterogeneity among studies, *e.g*. cell types and proportions. Alternatively, *mnnCorrect* utilizes mutual nearest neighbors (MNN) to account for heterogeneity among batches, recognizing matching cell types via MNN pairs [20]. By identifying the corresponding cells, a cell-specific correction can be learned for each MNN pair. As a consequence of local batch correction, *mnnCorrect* avoids the assumption of similar cell population compositions between batches assumed by previous methods. Following *mnnCorrect*, a series of new methods have been developed to integrate scRNA-seq data from different experiments [23–27]. For example, *Seurat v3* [23] uses MNN pairs between the reference batch and query batches to detect “anchors” in the reference batch. “Anchors” represent cells in a shared biological state across batches and are further used to guide the batch correction process through CCA. *BBKNN* [24] leverages neighborhood graphs to more efficiently cluster and visualize cell types. More recently, scRNA-seq batch correction is conducted by using deep learning approaches. For example, *scVI* [28] utilizes deep generative models to approximate the underlying distributions of the observed expression profiles, and can be used in multiple analysis tasks including batch correction. However, most existing batch correction methods for scRNA-seq data rely on similarities between individual cells, which do not fully utilize the clustering structures of different cell populations to identify the optimal batch-corrected subspace.

In this paper, by considering scRNA-seq data from different batches as different domains, we took advantage of the domain adaptation framework in deep transfer learning to properly remove batch effects by finding a low-dimensional representation of the data. The proposed method, *BERMUDA* (Batch-Effect ReMoval Using Deep Autoencoders), utilizes the similarities between cell clusters to align corresponding cell populations among different batches. We demonstrate that *BERMUDA* outperforms existing methods at combining different batches and separating cell types in the joint dataset based on UMAP visualizations and proposed evaluation metrics. By optimizing the maximum mean discrepancy (MMD) [29] between clusters across different batches, *BERMUDA* combines batches with *vastly different cell population compositions* as long as there is one common cell type shared between a pair of batches. Compared to existing methods, *BERMUDA* can also better preserve biological signals that exist in a subset of batches when removing batch effects. These improvements provide a novel deep learning solution to a persistent problem in scRNA-seq data analysis, while demonstrating state-of-the-art practice in batch-effect correction.

## Results

### Framework of *BERMUDA*

We propose *BERMUDA*, a novel unsupervised framework to remove batch effects across different batches by training an autoencoder (Figure 1a). After preprocessing the scRNA-seq data to select highly variable genes, we first used a graph-based clustering algorithm to detect cell clusters in each batch individually. Then, we applied a correlation-based approach to evaluate the similarity between cell clusters from different batches. Each pair of cell clusters was assigned a similarity score, which was later used as the coefficient in the transfer loss. Next, an autoencoder was trained using the standardized and scaled transcript-per-million (TPM) value of highly variable genes to learn a low-dimensional embedding of the gene expression profiles where the systematic biases across different batches were removed (Figure 1b). In order to successfully remove batch effects, we propose a novel approach by combining the reconstruction loss with the transfer loss when training the autoencoder. The reconstruction loss was calculated between the input and output of the autoencoder, which helped to learn a low-dimensional embedding that properly represented the original high-dimensional gene expression data. The transfer loss was calculated by estimating the difference of distributions between pairs of cell clusters using the lowdimensional representation, which helped to merge similar clusters from different batches. The *Mini-batch gradient descent* algorithm in deep learning was used to train *BERMUDA* where reconstruction loss and transfer loss were calculated from a sampled “mini-batch” during each iteration of the training process. The total loss in each iteration was then calculated by adding reconstruction loss and transfer loss with a regularization parameter (Equation 8) and the parameters in *BERMUDA* were then updated using gradient descent. Finally, the low-dimensional code learnt from the trained autoencoder was used for further downstream analysis.

**Figure 1.**
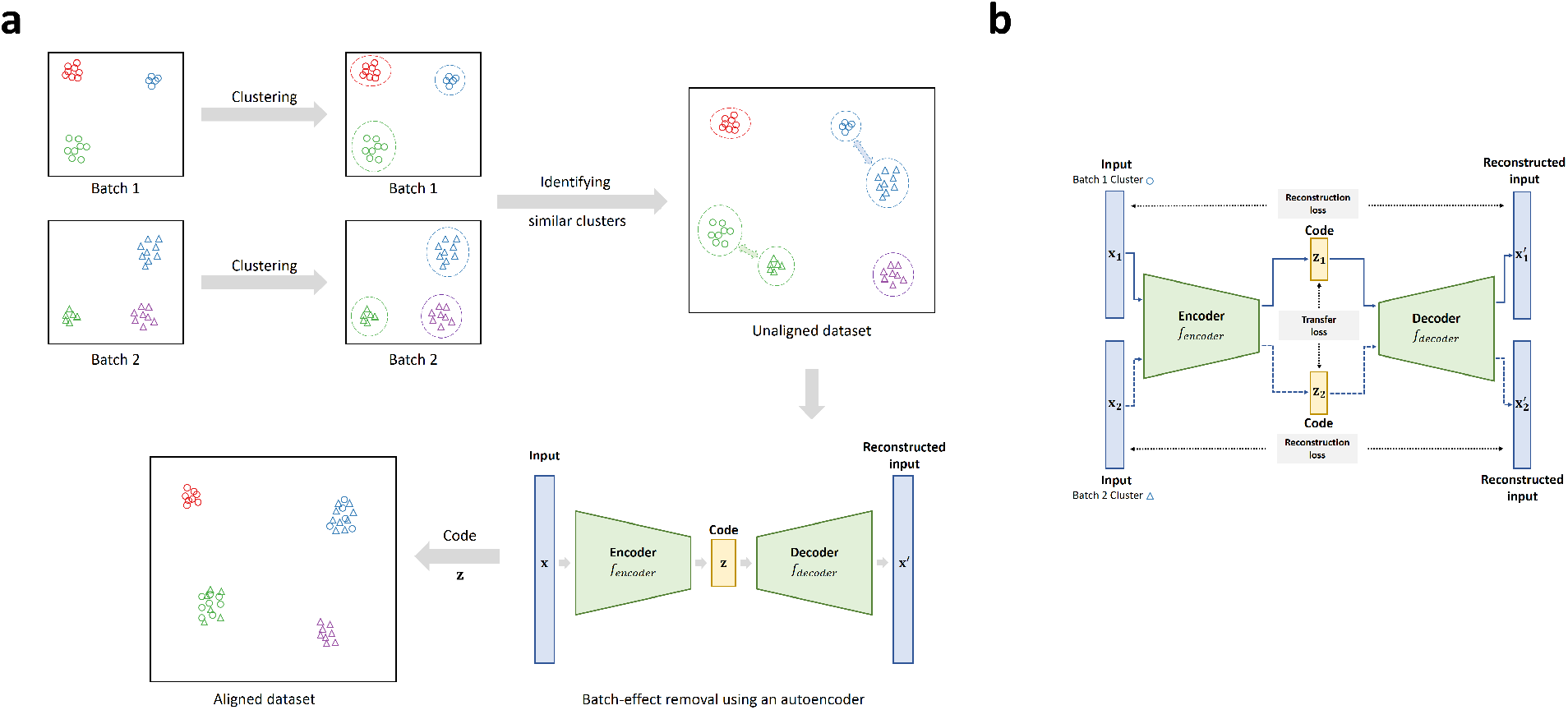
Overview of *BERMUDA* for removing batch effects in scRNA-seq data. a. The workflow of *BERMUDA*. Circles and triangles represent cells from *Batch 1* and *Batch 2*, respectively. Different colors represent different cell types. A graph-based clustering algorithm was first applied on each batch individually to detect cell clusters. Then, MetaNeighbor, a method based on Spearman correlation, was used to identify similar clusters between batches. An autoencoder was subsequently trained to perform batch-correction on the code of the autoencoder. The code of the autoencoder is a low-dimensional representation of the original data without batch effects and can be used for further analysis. b. Training an autoencoder to remove batch effects. The blue solid lines represent training with the cells in *Batch 1* and the blue dashed lines represent training with cells in *Batch 2*. The black dashed lines represent the calculation of losses. The loss function we optimized contains two components: the reconstruction loss between the input and the output of the autoencoder, and the MMD-based transfer loss between the codes of similar clusters.

The popularly used UMAP [30] method was utilized to visualize the cell type clustering results. In addition, three evaluation metrics are proposed to evaluate the performance of *BERMUDA: divergence_score, entropy_score, and silhouette_score* (See the “Methods” section). *Divergence_score* is an average of divergence of shared cell populations between pairs of batches, which indicates whether shared cell populations among different batches are mixed properly. *Entropy _score* is an average of local entropy of distinct cell populations between pairs of batches, which can evaluate whether cell populations not shared by all the batches remain separate from other cells after batch correction. *Silhouette_score* is calculated using cell type labels as cluster labels, which measures the quality of cell type assignment in the aligned dataset.

### Compare the performance of *BERMUDA* versus existing methods under different cell population compositions

We compared the performance of *BERMUDA* versus several existing state-of-the-art batch correction methods for scRNA-seq data (*mnnCorrect* [20], *BBKNN* [24], *Seurat v2* (v2.3.4) [22], *Seurat v3* (v3.0.0) [23], and *scVI* [28]) using four datasets (Table 1). To evaluate the performance of each method under different cell population compositions, we performed multiple data analysis experiments on each dataset with cell type labels (Table 2). For some experiments, we removed some cell types from specific batches to create different cell type distribution configurations. We performed three different experiments *(“Experiment all”, “Experiment removal1”*, and *“Experiment removal2”*) on each of the simulated datasets (2D Gaussian dataset and Splatter dataset). Specifically, for *Experiment all*, we applied each method to all the cells in the dataset. For *Experiment removal1*, we removed Type1 from *Batch1* and applied each method on this reduced dataset. For *Experiment removal2*, we removed Type1 from *Batch1* and Type4 from *Batch2* at the same time. We performed two different experiments (*“Experiment all”, “Experiment removal”*) on the human pancreas dataset. For *Experiment all*, we applied each method to all the cells in the entire dataset. For *Experiment removal*, we removed alpha and beta cells from *Baron batch* and alpha and beta cells from *Segerstolpe batch* (if applicable) and evaluated each method using this reduced dataset. We also applied *BERMUDA* to two batches of peripheral blood mononuclear cells (PBMCs).

**Table 1.** Datasets used for evaluation of *BERMUDA*.

| Dataset | Batch | Protocol | Number of cells | Cell Types (Number of cells) |
| --- | --- | --- | --- | --- |
| 2D Gaussian | Batch1 | Simulation | 2000 | Type1 (855), Type2 (237), Type3 (373), Type4 (535) |
|  | Batch2 | Simulation | 2000 | Type1 (379), Type2 (656), Type3 (102), Type4 (863) |
| Splatter | Batch1 | Splatter | 2000 | Type1 (810), Type2 (616), Type3 (388), Type4 (186) |
|  | Batch2 | Splatter | 1000 | Type1 (439), Type2 (262), Type3 (201), Type4 (98) |
| Pancreas | Muraro | CEL-Seq2 | 2042 | alpha (812), beta (448), gamma (101), delta (193), epsilon (3), acinar (219), ductal (245), endocrine (21) |
|  | Baron | inDrop | 8012 | alpha (2326), beta (2525), gamma (255), delta (601), epsilon (18), acinar (958), ductal (1077), endocrine (252) |
|  | Seegerstolpe | SMART-seq2 | 2061 | alpha (886), beta (270), gamma (197), delta (114), epsilon (7), acinar (185), ductal (386), endocrine (16) |
| PBMC | PBMC | 10X Chromium | 8381 |  |
|  | Pan T Cell | 10X Chromium | 3555 |  |

For the consistency in the paper, we refer to each pancreas and PBMC dataset, e.g., the Muraro dataset, as a batch, and refer to multiple batches investigating similar biological systems, e.g., pancreas, as a
dataset. Type1 to Type4 in the 2D Gaussian dataset and Splatter dataset refer to the virtual cell types that were generated using simulation techniques.

**Table 2.** Experiments performed for comparing *BERMUDA* with existing methods.

| Dataset | Batch | Experiment Name | Cell Type Removed |
| --- | --- | --- | --- |
| 2D<br>Gaussian | *Batch1, Batch2* | *Experiment all* | NA |
|  |  | *Experiment removal1* | Type1 from *Batch1* |
|  |  | *Experiment removal2* | Type1 from *Batch1*,<br>Type4 from *Batch2* |
| Splatter | *Batch1, Batch2* | *Experiment all* | NA |
|  |  | *Experiment removal1* | Type1 from *Batch1* |
|  |  | *Experiment removal2* | Type1 from *Batch1*,<br>Type4 from *Batch2* |
| Pancreas | *Muraro batch,*<br>*Baron batch* | *Experiment all* | NA |
|  |  | *Experiment removal* | alpha and beta from *Baron batch* |
|  | *Muraro batch,*<br>*Baron batch,*<br>*Seegerstolpe batch* | *Experiment all* | NA |
|  |  | *Experiment removal* | alpha and beta from *Baron batch*,<br>alpha and beta from *Seegerstolpe batch* |
| PBMC | *PBMC batch,*<br>*pan T cell batch* | NA | NA |

### *BERMUDA* outperformed existing methods in removing batch effects on simulated data

To assess the performance of *BERMUDA* for batch-effect removal, we first applied it to a simulated dataset (referred to as “2D Gaussian dataset”) with four shared virtual cell types, where the expression profiles were generated from a 2-dimensional biological subspace following the method in [20] (See the “Methods” section). In order to recover and better visualize the underlying biological subspace, we set the number of neurons in the bottleneck layer of the autoencoder to two. We compared the performance of *BERMUDA* to other existing methods in three different scenarios:

- all the cells from the two batches (referred to as *“Experiment all”*);
- removing Type1 in *Batch1* (referred to as *“Experiment removal1”*);
- removing Type1 in *Batch1* and Type4 in *Batch2* at the same time (referred to as *“Experiment removal2”*).

*Experiment removal2* represented the most difficult case in these three scenarios because only two cell types were shared by both batches. We evaluated the results by inspecting the 2-dimensional visualizations (Figure 2a, Additional file 1: Figure S1). Ideally, the visualizations generated after proper batch correction should contain four separate cell clusters (each representing a cell type), and the clusters for shared cell types should contain a homogenous mixture of cells from both batches. In *Experiment removal2, BERMUDA* properly removed batch effects (Figure 2a1-2) while both *mnnCorrect* and *BBKNN* generated false similarities between cell types that did not exist in the original data (Figure 2a4-5). Specifically, in *Experiment removal2*, clusters corresponding to Type1 and Type4 were closely connected in the *mnnCorrect* results (Figure 2a4), and clusters corresponding to Type1, Type3, and Type4 were not as well separated in the *BBKNN* results (Figure 2a5). Moreover, only *BERMUDA* produced proper batch correction consistently across three different experiments (Figure 2a, Additional file 1: Figure S1). For example, *BBKNN* incorrectly separated Type1 from the same batch into two distinct clusters in *Experiment removal1*(Additional file 1: Figure S1b5). This suggests that although *mnnCorrect* and *BBKNN* can handle differences in cell population composition among different batches, their performance was less optimal compared with *BERMUDA* when such differences were large.

**Figure 2.**
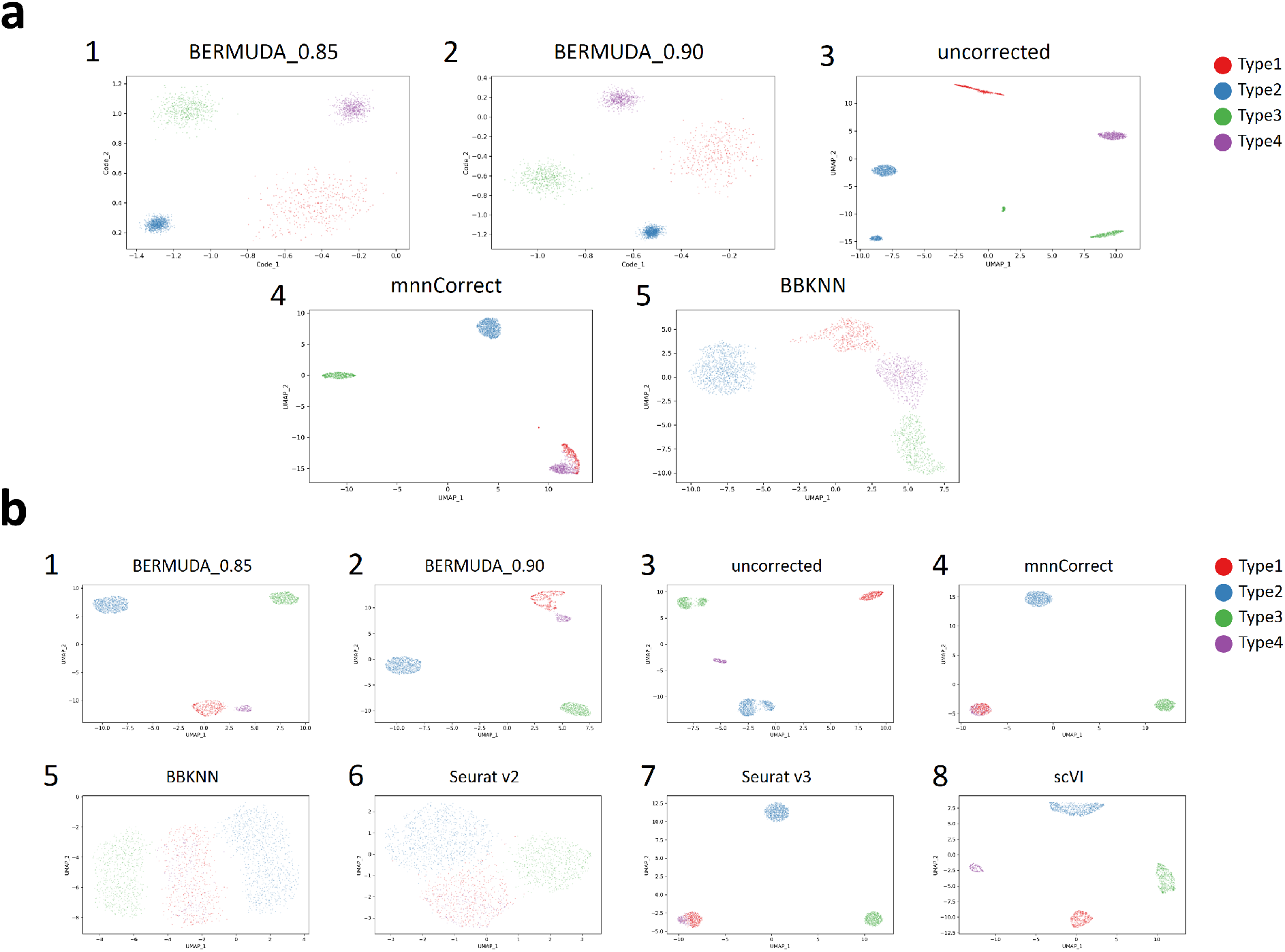
Removing batch effects in simulated scRNA-seq data. a. UMAP visualizations of results for *Experiment removal2* on 2D Gaussian dataset, where Type1 from *Batch1* and Type4 from *Batch2* were removed. BERMUDA_0.85 and BERMUDA_0.90 represents results of *BERMUDA* with *S_thr_* = 0.85 and 0.90, respectively. b. UMAP visualizations of results for *Experiment removal2* on Splatter dataset, where Type1 from *Batch1* and Type4 from *Batch2* were removed.

In order to compare *BERMUDA* with the popularly used scRNA-seq data analysis tool *Seurat v2* [22] and the recently-proposed *Seurat v3* [23] and *scVI* [28] (where the workflows were designed for scRNA-seq count data), we evaluated *BERMUDA* using two batches of simulated single-cell RNA sequence counts generated by Splatter [31] (referred to as “Splatter dataset”). We conducted multiple batch correction experiments using different cell populations:

- all the cells from two batches (referred to as *“Experiment all”*);
- removing Type1 in *Batch1* (referred to as *“Experiment removal1”*);
- removing Type1 in *Batch1* and Type4 in *Batch2* (referred to as *“Experiment removal2”*).

Again, *Experiment removal2* was the most difficult scenario. When only two cell types were shared between two batches in *Experiment removal2*, only *BERMUDA* properly removed batch effects (Figure 2b1-2). Although *scVI* could also align corresponding cell types (Figure 2b8), it failed to merge cells from different batches within each cell type (Additional file 1: Figure S2c8). The other methods improperly merged Type1 with Type4 in the UMAP visualizations (Figure 2b4-7). Moreover, *BERMUDA* was the only method that could consistently remove batch effects in all three cases. When all four cell types were shared in both batches, we observed that all the methods properly merged cells of the same type (Additional file 1: Figure S2a). However, *BBKNN* and *Seurat v2* produced much lower *silhouette_score* values due to the inflated variance within cell clusters compared to the original data (Additional file 1: Figure S2d1). Also, when the difference in cell type distributions was introduced in *Experiment removal1*, we observed that *BBKNN* and *Seurat v2* could no longer mix the same cell type from different batches properly (Additional file 1: Figure S2b5-6). The results of *Seurat v2* were anticipated since it was designed to align different batches globally without considering the population differences among batches. The inconsistent results of *BBKNN* under different cell type distributions indicated that the neighborhood graphs generated by *BBKNN* may not always be reliable due to the fact that the method mainly focused on computational efficiency.

The observations through UMAP visualizations were further confirmed by evaluating the results through the proposed *divergence_score*, *entropy_score*, and *silhouette_score* (Additional file 1: Figure S2d). In *Experiment removal2, scVI* could not properly align cells from different batches within each cell type, resulting in a high *divergence_score* (Additional file 1: Figure S2d5-6). Other existing methods produced low *silhouette_score* values (Additional file 1: Figure S2d6-7) since they were not able to sperate different cell types correctly. We observed that *BERMUDA* consistently yielded the best performance in all three cases when evaluated using the proposed metrics (Additional file 1: Figure S2d).

By evaluating *BERMUDA* on two different simulated datasets, we demonstrate that it achieves better performance than existing methods in removing batch effects, especially when the difference of cell population compositions among batches is large.

### *BERMUDA* outperformed existing methods in removing batch effects on human pancreas data

To further evaluate *BERMUDA* using biological data, we applied it to publicly available human pancreas datasets that were generated utilizing different scRNA-seq protocols. Muraro *et al*. [32] used CEL-Seq2, a multiplexed linear amplification RNA sequencing technique. This dataset will be referred to as *“Muraro batch.”* Baron *et al*. [33] used a droplet RNA-seq technology. We refer to this dataset as *“Baron batch.”* In addition to evaluating our method using all the cells in both batches (referred to as *“Experiment all”*, Figure 3a, Additional file 1: Figure S4a), we also removed alpha and beta cells from *Baron batch* (referred to as *“Experiment removal”*, Additional file 1: Figure S4b-c) to simulate vast differences in cell type distributions found in real scRNA-seq data.

**Figure 3.**
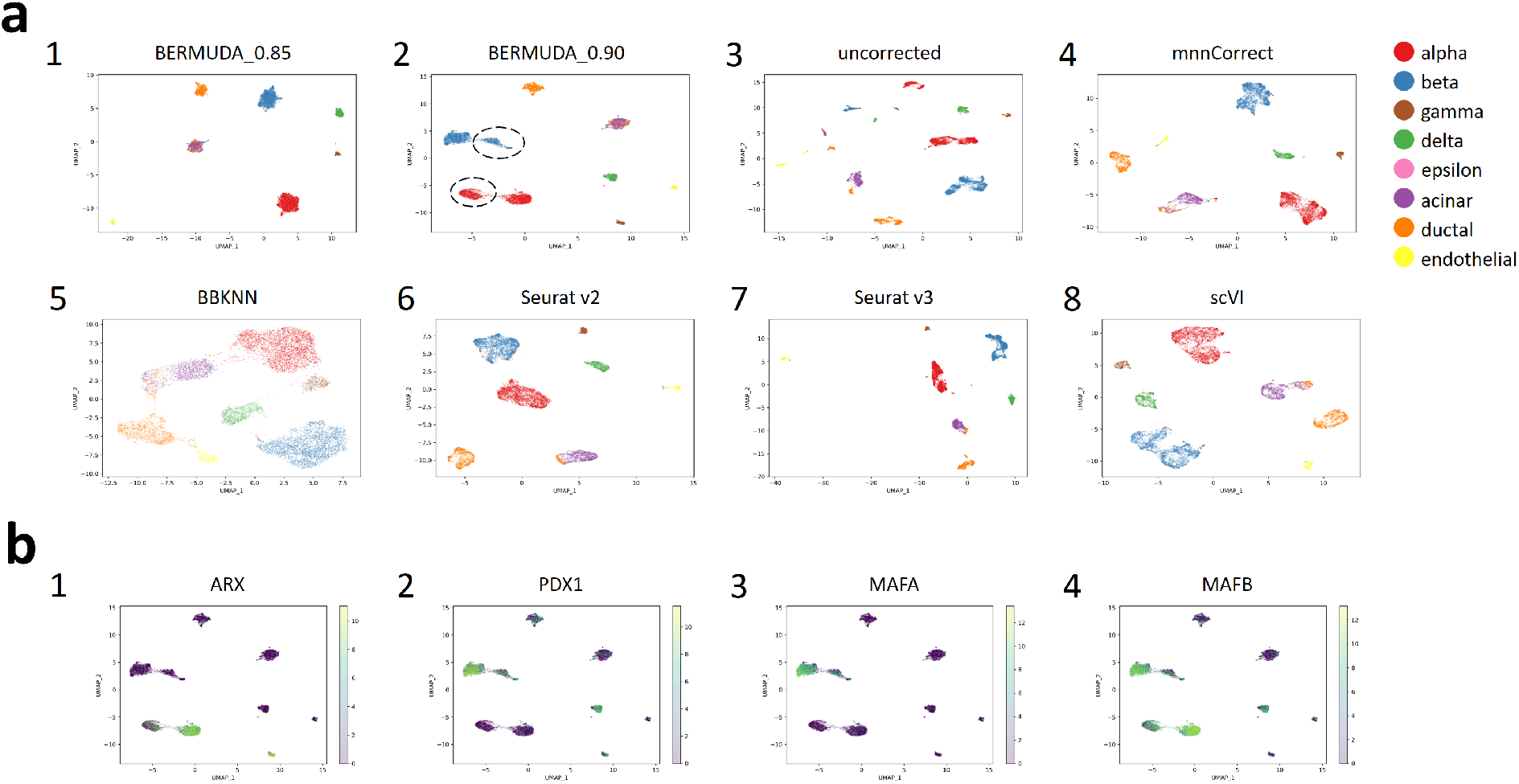
Removing batch effects in scRNA-seq data of pancreas cells. a. UMAP visualizations of batch-effect removal results for *Experiment all* on pancreas dataset. Identified alpha and beta cell subpopulations in the *Baron batch* are highlighted with dashed circles. b. Expression patterns of differently expressed genes within alpha and beta cells colored by log-transformed TPM values. Statistical significance of differential expression analysis is listed in Table S1-S2.

*BERMUDA* achieved competitive results in both cases compared to *mnnCorrect, BBKNN, Seurat v2, Seurat v3*, and *scVI*. Specifically, in *Experiment removal*, the performance of *Seurat v2* deteriorated when alpha and beta cells were removed from the *Baron batch* (Additional file 1: Figure S4d) because only utilizing CCA was not capable of addressing the case where only a subset of cell types was shared among batches. *BBKNN* and *scVI* separated different cell types into different clusters, but failed to mix cells from different batches within clusters properly (Figure 3a5, 3a8, Additional file 1: Figure S4a5, S4a8), producing high *d?vergence_score* values (Additional file 1: Figure S4d1). *Seurat v3* and *mnnCorrect* were designed to cope with cell type distributions across datasets being different. However, *Seurat v3* could not produce consistent results with clear separation between cell types when the cell population composition varied within the same set of data. For example, *Seurat v3* produced a UMAP visualization that represented alpha, beta, gamma, and delta cells as tightly connected clusters when alpha and beta cells from the *Baron batch*were removed. This differed from both the original data and the *Seurat v3* result in *Experiment all* (Figure 3a7, Additional file 1: Figure S4b7). The *mnnCorrect* method produced relatively good results across both cases (Figure 3a4, Additional file 1: Figure S4b4). However, *BERMUDA* still outperformed all of these existing methods when evaluated using the three proposed metrics simultaneously (Additional file 1: Figure S4d).

### *BERMUDA* preserved batch-specific biological signals in human pancreas data

In this paper, we use “batch-specific” biological signals to refer to *bona fide* biological signals that are not shared among all the batches. A homogeneous mixture of the same cell type among different batches is an important indicator of proper batch corrections. However, an overly batch-corrected homogeneous mixture could lead to loss of the subtle batch-specific biological signals that may contain information about potential biomarkers of batch-specific cell subtypes, which could defeat the purpose of carrying out scRNA-seq experiments. One advantage of *BERMUDA* is the ability to balance between maintaining a more homogeneous mixture of cell types and retaining the subtle structures within cell types, which is accomplished by adjusting the value of *S_thr_*. A lower *S_thr_* value can merge the same cell type from different batches more homogeneously. However, increasing the value of *S_thr_* can also be helpful, as it allows subtle batch-specific biological signals to be preserved. We demonstrate that *S_thr_* values between 0.85 and 0.90 can produce proper batch correction in different datasets consistently (Additional file 1: Figure S8-S9). For example, in *Experiment all* on the pancreas dataset, *BERMUDA* produced competitive results for both *S_thr_*values (Additional file 1: Figure S4d1). Each cell type was separated as a single cluster with a homogenous mixture of cells from both batches when *S_thr_* = 0.85 (Figure 3a1). In addition, we observed two closely connected cell clusters of alpha and beta cells from the *Baron batch* when *S_thr_* = 0.90 (Figure 3a2), which was consistent with the original, uncorrected data (Figure 3a3). When *S_thr_* = 0.90, only one of two connected clusters in alpha or beta cells in *Baron batch* merged with the corresponding cells in the *Muraro batch*, whereas the other remained unmixed (Figure 3a2, Additional file 1: Figure S4a2). This observation was confirmed by the *mnnCorrect* and *Seurat v3* results, although both with a subtler distinction of clusters (Figure 3a4, 3a7). The consistent observation indicated that observed cell clusters inferred alpha or beta cell subtypes that only appeared in the *Baron batch*, which could be detected when *S_thr_* = 0.90.

To further investigate the potential cell subtypes, we performed differential gene expression analysis within alpha cells and beta cells on two pairs of cell populations. For alpha cells, we examined differential gene expression between the mixed cluster from *Muraro batch* versus the unmixed cluster from *Baron batch*. We also separately examined differential gene expression between the mixed and unmixed cluster from *Baron batch*. For beta cells, we performed the same two sets of differential gene expression analysis between the respective mixed cluster and the unmixed cluster. We used the “FindMarkers” function in *Seurat v2* [22] to identify genes that have significantly different expression patterns in both pairs of cell populations (Additional file 1: Table S1-S2), which resulted in genes related to important pancreatic functions. For example, *ARX* and *MAFB* (Table S1, Figure 3b1, 3b4) were significantly under-expressed in the unmixed alpha cells compared with the mixed alpha cells. *PDX1, MAFA*,and *MAFB* were significantly under-expressed in the unmixed beta cells compared with the mixed beta cells (Table S2, Figure 3b2-4). Previous studies [34, 35] have shown that *ARX* plays a key role in the differentiation of pancreatic islet cells. Moreover, the decrease of *MAFA, MAFB*, and *PDX1* expression levels have been found in human Type 2 diabetes islet alpha and beta cells, which can be associated to islet cell dysfunction [36]. Identification of the aforementioned transcriptomic signatures suggested that the refined cell clusters that only exist in *Baron batch* may contain cell subpopulations with altered pancreatic functions, which may warrant further biological investigation. As mentioned above, such cell subpopulations were also identified in *Seurat v3* and *mnnCorrect*.

By applying our method to human pancreas datasets, we demonstrate that *BERMUDA* effectively removes batch effects under different cell population compositions across batches. *BERMUDA* outperforms existing methods in combining corresponding cell types, preserving subtle cell clusters that are not shared by all the batches, and properly separating different cell types. We also show that *BERMUDA* can preserve and even amplify biologically meaningful structures within cell types when integrating different batches.

### *BERMUDA* properly mapped human pan T cells to PBMCs

In order to show that *BERMUDA* can transfer information between batches with more complicated cell populations and reveal biological signals that might remain hidden when analyzing each batch individually, we also applied *BERMUDA* to two large scRNA-seq datasets of peripheral blood mononuclear cells (PBMCs) generated from the 10X Genomics Chromium platform *(i.e., “PBMC batch”* and *“pan T cell batch”). PBMC batch* and *pan T cell batch* were acquired from different healthy donors. We expected that cell types in the *pan T cell batch* should be a subset of those in the *PBMC batch*, while the *PBMC batch* could contain other cell types such as B cells and monocytes. From the UMAP visualization of the uncorrected data (Additional file 1: Figure S5c), we speculated that the *pan T cell batch* should roughly correspond to the largest cell cluster in the *PBMC batch*, while the other two smaller clusters should represent non-T cells in the *PBMC batch*. As shown in Figure 4b, *BERMUDA* successfully retained the two PBMC-specific clusters, while mapping the T cell population properly.

**Figure 4.**
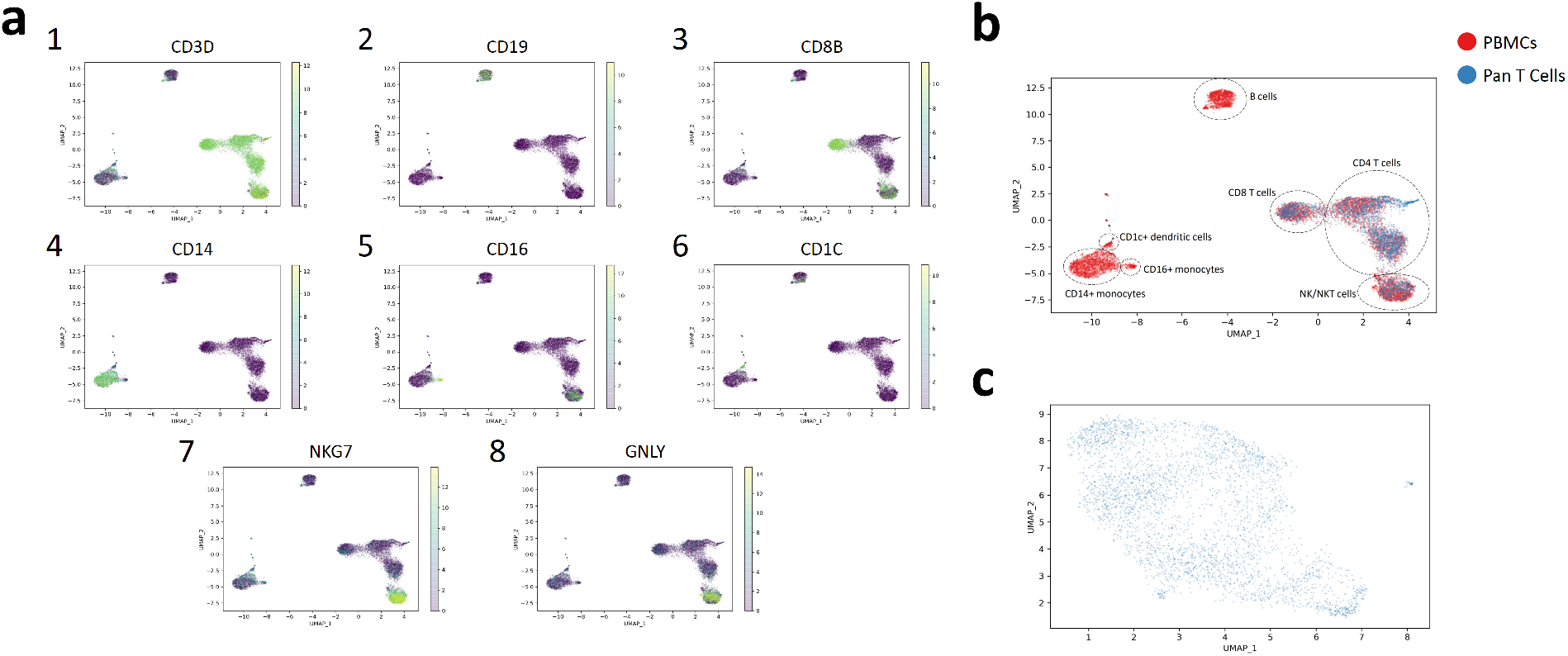
Removing batch effects in scRNA-seq data of PBMCs. a. Expression patterns of marker genes of immune cells colored by log-transformed TPM values. b. The UMAP visualization of results produced by *BERMUDA* colored by batches. Different cell types were identified by analyzing expression patterns of maker genes and are highlighted by dashed circles. *BERMUDA* correctly combined the corresponding cell types between different batches while preserved cell types not shared by both batches as separate clusters. c. The UMAP visualization of *pan T cell batch*. No obvious clustering structure was observed when visualizing *pan T cell batch* individually.

Since the obtained data did not include pre-defined cell type labels, we evaluated the results by inspecting the expression patterns of marker genes using the UMAP visualizations. We identified seven major cell types in our results through corresponding marker genes: CD4 T cells (Figure 4a1, 4a3), CD8 T cells (Figure 4a3), B cells (Figure 4a2), CD1c+ dendritic cells (Figure 4a6), CD14+ monocytes (Figure 4a4), CD16+ monocytes (Figure 4a5), and NK/NKT cells (Figure 4a7-8). Cell types that only existed in the *PBMC batch* (B cells, CD14+ monocytes, CD16+ monocytes, CD1c+ dendritic cells) retained a similar structure with the uncorrected original data (Additional file 1: Figure S5a-c), which further demonstrated that our method can combine batches while preserving biological information that only existed in a subset of batches. Interestingly, the UMAP visualization of *pan T cell batch* alone did not show any clear clustering structures. However, different cell types within *pan T cell batch* were further distinguished as different clusters by combining with PBMC cells. This indicates that *BERMUDA* is capable of effectively transferring the biological information from one batch to another and extracting biological insights from batches that might not be evident by studying each batch individually.

Conversely, *Seurat v2* and *Seurat v3* failed to align the overlapping cell types from two batches properly (Additional file 1: Figure S5f-g). *BBKNN* overly corrected batch effects and lost the structure within the T cell cluster and the monocyte cluster observed in the original uncorrected data (Additional file 1: Figure S5e). *mnnCorrect* and *scVI* were able to align the shared cell types between two batches. However, *scVI* did not produce a homogeneous mixture of both batches within pan T cells, while *mnnCorrect* produced a more heterogeneous structure within the T cell cluster when comparing to the uncorrected data visually (Additional file 1: Figure S5d, S5h).

### *BERMUDA* can be generalized to combine multiple batches

Since *BERMUDA* is based on applying MMD loss to similar clusters from different batches for batch-effect removal, it can be easily generalized to deal with multiple batches simultaneously. Here we demonstrate such capability by including another human pancreas dataset generated by Segerstolpe *et al*.[37] (referred to as *“Segerstolpe batch*”). The *Segerstolpe batch* was generated using the sequencing platform SMART-seq2 technology and contained 2,061 cells of interest.

Similar to the previous scenarios, two experiments were performed on the human pancreas dataset: one with all the cells in the dataset (referred to as *“Experiment all”*, Additional file 1: Figure S6) and one with alpha and beta cells removed from both the *Baron batch* and the *Segerstolpe batch* (referred to as *“Experiment removal’*, Additional file 1: Figure S7). *BERMUDA* consistently produced competitive results in both experiments (Additional file 1: Figure S6c, S7c). Moreover, the aforementioned potential alpha and beta cell subpopulations in the *Baron batch* were also consistently observed when setting *S_thr_* = 0.90. This demonstrates that our method can naturally handle multiple batches simultaneously and generate consistent and robust results when number of batches increases.

## Discussion

With the rapid advances of single cell sequencing technologies and the accumulation of large scRNA-seq data, it is important to combine data from different samples or studies to fully harness the power of scRNA-seq techniques, infer cell lineages, and identify *bona fide* transcriptional signals. Proper removal of batch effects between scRNA-seq experiments has become an urgent issue. Batch effects arise from many potential sources, such as different protocol-, platform-, or lab-specific artifacts. In addition, cell type compositions differ among scRNA-seq experiments because of underlying biological variation. This increases the difficulty of distinguishing *bona fide* biological signals from systematic biases and accurately representing combined data from multiple sources.

In this study, we propose *BERMUDA*, a novel batch correction method based on the deep transfer learning framework. We considered scRNA-seq data from different batches as different domains and utilized domain adaptation approaches to project the original data to a lower-dimensional space without batch effects. We demonstrated that *BERMUDA* can properly combine data from different batches in both simulated data and real datasets as long as there is one common cell type shared by a pair of batches. We observed that *BERMUDA* outperformed existing methods *(mnnCorrect, BBKNN, Seurat v2, Seurat v3* and *scVI)* in three aspects: homogeneously mixing the same cell type from different batches, maintaining the purity of batch-specific cell clusters, and separating different cell types in the combined dataset. By training an autoencoder with MMD loss on pairs of similar clusters, *BERMUDA* can effectively perform batch correction without considering the specific sources of batch effects. Because of this, *BERMUDA* is capable of removing batch effects resulting from various sources, such as simulated batch effects, batch effects from the same platform in the PMBC dataset, and batch effects generated by different platforms in the human pancreas dataset. Moreover, by using a deep neural network framework and training through the mini-batch gradient descent algorithm, *BERMUDA* is scalable to large numbers of cells, which is beneficial for the rapid adoption of scRNA-seq experiments.

*BERMUDA* only requires batches to share one common cell type with another batch, which is distinct from existing methods that do not account for cell type distributions across batches. Because *BERMUDA* performs batch correction based on similarity among cell clusters, it accommodates variation in both cell composition across batches and also different cell populations. Rather than considering a global batch effect, *BERMUDA* removes batch effects locally by combining similar clusters while maintaining the global structure of the data. By applying this approach, knowledge from one batch can be transferred to another, which augments biological insights that cannot be observed when examining each batch individually. For example, when directly visualizing the *pan T cell batch*, it did not show obvious clustering structures (Figure 4c). However, by combining it with PBMCs, pan T cells were properly mapped to the corresponding cell clusters in the *PBMC batch*, providing a higher resolution of cell subtypes within the *pan T cell batch* (Figure 4b).

Typically, there is a trade-off between removing batch effects and retaining experimental specific biological signals, causing some existing methods (such as *BBKNN* and *Seurat v2)* to lose subtle biological signals while merging multiple batches for a more homogenous mixture of cells [38, 39]. *BERMUDA* can preserve and amplify the sensitivity to biological differences in the original data while properly removing batch effects. Specifically, the trade-off between a more homogeneous mixture within cell types and preserving more batch-specific biological signals could be adjusted by changing the *S_thr_* value, where a higher *S_thr_* value can help to retain more biological signals that only exist in a subset of batches. We demonstrate experimentally that *BERMUDA* is robust and can outperform existing methods in a wide range of *S_thr_* values, and choosing *S_thr_* between 0.85 and 0.90 can consistently yield a good balance between proper batch correction and preserving batch-specific biological signals (Additional file 1: Figure S8, S9).

*BERMUDA* also has limitations. Since the design of *BERMUDA* is based on similarity between cell clusters, it can remove batch effects from scRNA-seq data with distinct cell populations effectively. However, clusters may not always be well separated due to technical or biological noise. Moreover, scRNA-seq data can also be continuously variable — such as data generated for cell differentiation. While *BERMUDA* was originally designed with a focus on scRNA-seq data with distinct cell populations, it can also accommodate such data by adjusting the resolution in the graph-based clustering algorithm and the trade-off between reconstruction loss and transfer loss to align clusters at a more granular level. It is also the focus of our future work to improve *BERMUDA* to accommodate these data even more effectively. Another limitation is that the use of k-nearest neighbor in the clustering algorithm (integrated in *Seurat v2)* may not scale well to extremely large datasets [24]; though, a neural-network-based framework for batch correction is capable of accommodating large datasets. However, novel clustering algorithms could be applied in the future to speed up the clustering process. For example, Ren *et al*. [40] proposed a novel framework for clustering large scRNA-seq data, which reduced the computational complexity to O(n) while maintaining a high clustering accuracy.

## Conclusions

Removing batch effects is essential for the analysis of data from multiple scRNA-seq experiments and multiple technical platforms. Here we introduce *BERMUDA*, a novel batch-correction method for scRNA-seq data based on deep transfer learning. We use an autoencoder to learn a low-dimensional representation of the original gene expression profiles while removing the batch effects locally by incorporating MMD loss on similar cell clusters. *BERMUDA* provides several improvements compared to existing methods. Firstly, by introducing three different metrics to evaluate the batch correction performance we demonstrate that *BERMUDA* outperforms existing methods in merging the same cell types, preserving cell types not shared by all the batches, and separating different cell types. Secondly, *BERMUDA* can properly remove batch effects even when the cell population compositions across different batches are vastly different. Thirdly, *BERMUDA* can preserve batch-specific biological signals and discover new information that might be hard to extract by analyzing each batch individually. Finally, *BERMUDA* can be easily generalized to handle multiple batches and can scale to large datasets.

## Methods

### Datasets used for performance evaluation

For consistency, in this paper an individual dataset will be referred to as a “batch.” Multiple batches investigating similar biological problems will be referred to as a “dataset.” We applied *BERMUDA* on simulated datasets, human pancreas cell datasets, and PBMC datasets to assess its performance (Table 1). We used two methods to generate simulated datasets for evaluating the performance of *BERMUDA*. For the 2D Gaussian dataset, we followed [20] to generate highly synthetic data with batch effects. We simulated two batches of four different cell types according to different bivariate normal distributions in a 2-dimensional biological subspace. The cell population composition of each batch was generated randomly. Then we randomly projected the data to a 100-dimensional space to simulate the high-dimensional gene expression data. Gene-specific batch effects were generated by adding Gaussian noise to the highdimensional data. For the Splatter dataset, we used the Splatter [31] package to simulate RNA sequence counts of two batches with four different cell populations. Splatter can directly simulate multiple batches following similar cell type compositions at the same time. We set the cell population composition to be 0.4, 0.3, 0.2, and 0.1 among the four simulated cell types.

To evaluate whether *BERMUDA* can remove batch effects in real scRNA-seq data and extract meaningful biological insights, we also applied it to datasets of human pancreas cells and PBMCs. The pancreas dataset was obtained from Gene Expression Omnibus (GSE85241 for *Muraro batch* [32], GSE84133 for *Baron batch* [33]) and The European Bioinformatics Institute (E-MTAB-5061 for *Segerstolpe batch* [37]). The PBMC dataset was obtained from 10X Genomics support datasets. To effectively compare the difference between the cases where all the cell types or only a subset of those were shared among batches, we only retained the shared cell types in the pancreas dataset. The details of the datasets are shown in Table 1.

### Framework of *BERMUDA*

*BERMUDA* is a novel unsupervised framework for batch correction across different batches (Figure 1a). The workflow of *BERMUDA* includes the following five steps: 1. Preprocessing of scRNA-seq data. 2. Clustering of cells in each batch individually. 3. Identifying similar cell clusters across different batches. 4. Removing batch effects by training an autoencoder (Figure 1b). 5. Utilizing the batch-corrected codes for further downstream analysis. We introduce each step in detail in the following sections.

### Preprocessing

Gene expression levels from each cell were first quantified using transcript-per-million values (TPM). First, we restricted analysis to genes that were highly variable based on *Seurat v2* [22]. Gene expression values were normalized as

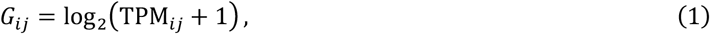

where TPM*_ij_* is the TPM of gene *i* in cell *j*. We subsequently standardized expression per batch as

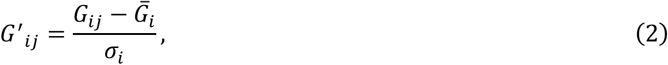

where *G′_ij_* is the standardized expression level of gene *i* in cell *j*. 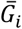 is the mean expression level for gene *i* and *σ_i_* is the standard deviation of expression level for gene *i*. Then, we linearly scaled the expression level of each gene to [0, 1].

### Clustering of cells and identifying similar cell clusters

Cell clusters were identified from each batch individually following the pipeline in *Seurat v2* [22]. *Seurat v2* implemented a clustering algorithm based on optimizing the modularity function on a k-nearest neighbor graph. We then used MetaNeighbor [8] to determine the similarity between clusters from different batches based on Spearman correlation. For *n* batches each contains *c_i_* clusters, *i* = 1,2, …, *n*, MetaNeighbor produces a similarity score for each pair of cell clusters by calculating the mean of area under the receiver operator characteristic curve (AUROC) in cross-validation. We denote *M*_*i*_1_,*j*_1_,*i*_2_,*j*_2__ as the similarity score between cluster *j*_1_ in batch *i*_1_ and cluster *j*_2_ in batch *i*_2_. Because we were interested in similar clusters across different batches, we set the similarity score between clusters within the same batch to 0. For each cluster, we considered the most similar cluster in each of the other batches, such that

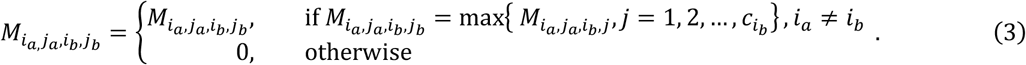

To accommodate the case where a cell cluster in one batch corresponds to multiple clusters in another batch and make *BERMUDA* more robust to the results in the clustering step, we made the similarity scores between two clusters symmetrical by modifying *M_i_a_,j_a_,i_b_,j_b__* as

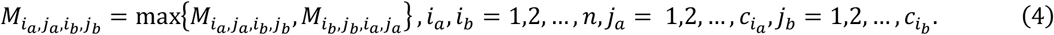

Finally, we binarized *M_i_a_,j_a_,i_b_,j_b__* with a threshold value *S_thr_*, where

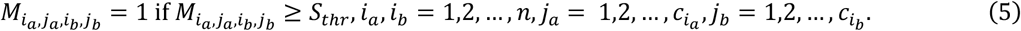

*S_thr_* was chosen empirically and we observed that *BERMUDA* achieves robust and competitive results across different datasets when *S_thr_* is between 0.85 and 0.90 (Additional file 1: Figure S8, S9).

### Batch correction using an autoencoder

*BERMUDA* uses an autoencoder to project the original uncorrected gene expression profiles to a lowdimensional space to remove the experimental artifacts across different batches (Figure 1b). An autoencoder can be represented as a function **x**′ = *f*(**x**) = *f_decoder_*(*f_encoder_*(**x**)), where *f* reconstructs the input gene expression profile **x** through the neural network. To avoid trivial solutions, autoencoders usually incorporate a bottleneck layer that learns a low-dimensional embedding of the input data called code, *e.g*., **z** = *f_encoder_*(**x**). In *BERMUDA*, we used an autoencoder with three hidden layers, and the default number of neurons in each hidden layer were 200, 20, 200. For the synthetic data generated from bivariate Gaussian distributions, we set the neurons in each hidden layer as 20, 2, 20 to reconstruct the 2-dimensional biological plane.

The mini-batch gradient descent algorithm commonly adopted in deep learning was used to train *BERMUDA*. Mini-batch gradient descent is a variation of the gradient descent algorithm and is widely adopted in the field of deep learning. For each iteration in each epoch during the training process of *BERMUDA*, a “mini-batch” **X** = {**x**_1_,**x**_2_, …, **x**_B_} was sampled from the dataset, which contained *n_mb_* cells from each cluster 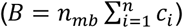. We used *n_mb_=* 50 in our experiments. However, we observed that *BERMUDA* was robust to the choice of *n_mb_* and can outperform existing methods under a wide range of *n_mb_* values (Additional file 1: Figure S10). The loss was calculated on the entire mini-batch and the parameters in *BERMUDA* were then updated using gradient descent. In each epoch, multiple iterations were performed to cover all the cells in the dataset.

The loss function for training the autoencoder consisted of two parts. The first part is a reconstruction loss between the output layer and the input layer defined by mean squared error

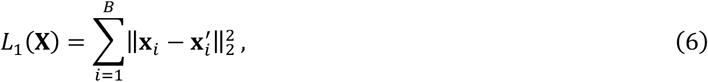

where **x**_*i*_ and 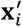 are input and reconstructed expression profile of the *i*-th cell in a mini-batch. The second part is a novel maximum mean discrepancy (MMD) based loss [29] that estimates the differences in distributions among similar cell clusters in different batches. MMD is a non-parametric distance estimate between distributions based on the reproducing kernel Hilbert space (RKHS) and has proven to be highly effective in many deep transfer learning tasks [41–44]. Since MMD does not require density estimates as an intermediate step and does not assume any parametric density on the data, it can be applied to different domains [45]. MMD is also memory-efficient, fast to compute, and performs well on high dimensional data with low sample size [46, 47]. Considering the case where only a subset of the cell population is shared among batches, instead of applying MMD loss on batches entirely, we only considered the loss between pairs of similar cell clusters among different batches. So, the MMD-based loss can be defined as

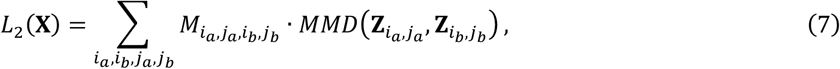

where **Z**_*i,j*_ is the code of the input **X**_*i,j*_, and **X**_*i,j*_ is the expression profiles of cells from cluster *j* of batch *i* in the mini-batch **X**. *MMD*(·) equals to zero when the underlying distributions of the observed samples are the same. By minimizing the MMD loss between the distributions of similar clusters, the autoencoder can be trained to remove batch effects in the bottleneck layer. In summary, the total loss function on a mini-batch can be written as

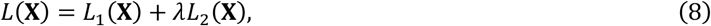

where *λ* is a regularization parameter. We followed the strategy introduced by Ganin *et al*. [48] to gradually increase *λ* during the training process. The regularization parameter at epoch *p* is calculated as

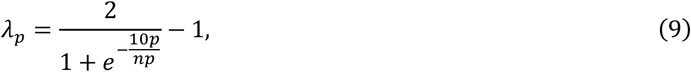

where *np* is the number of total epochs in training. This can help the autoencoder to first focus on finding a proper low-dimensional representation of the original gene expression data, then focus on aligning the distributions of similar clusters in the low-dimensional space.

### Performance evaluation

To evaluate the performance of *BERMUDA*, we examined the outputs when specific cell types were removed from their respective batches. We then used three metrics to compare algorithm performance. First, we used a k-nearest-neighbor based divergence estimation method [49] to evaluate the quality of merging the shared cell population among batches. For *N* scRNA-seq batches with gene expression profiles **X**_1_,**X**_2_, …, **X**_*N*_ and their corresponding batch-corrected low-dimensional embeddings **Z**_1_,**Z**_2_, …, **Z**_*N*_, we define

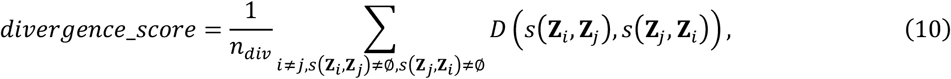

where *s*(**Z**_*i*_,**Z**_*j*_) is the cell population in **Z**_*i*_ that is shared by **Z**_*j*_, *D*(**Z**_*i*_,**Z**_*j*_) is the divergence estimation of the two distributions given samples **Z**_*i*_ and **Z**_*j*_, and 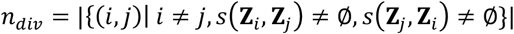 is the number of pairs of batches with shared cell population. Since proper removal of batch effects should produce results where the distributions of shared cell populations among different batches are similar, a smaller *divergence_score* is preferred, indicating that the shared cell population between different batches are homogeneously mixed. Second, we used entropy to evaluate whether a cell population that only exists in a certain batch remains distinct from other populations after correction. We define

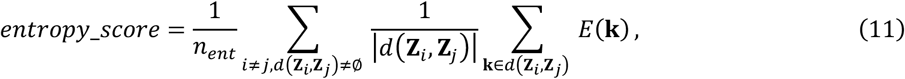

where *d*(**Z**_*i*_,**Z**_*j*_) is the cell population in **Z**_*i*_ that is not shared by **Z**_*j*_, and 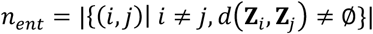 is the number of pairs of batches where there exists distinct cell population in **Z**_*i*_ from **Z**_*j*_. *E*(**k**) is the estimation of entropy locally around cell **k** defined as

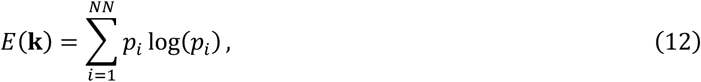

where *p_i_* is the proportion of cells from batch *i* among the *NN*-nearest neighbors of cell **k**. We chose *NN* = 100 in our evaluations. When batch effects are removed properly, a cell type that only exists in a batch should not be mixed with cells from other batches. So, a smaller *entropy_score* is desired, suggesting that biological signals only contained in a subset of batches are properly preserved during correction. Note that when all the batches share the same cell types, we did *not* calculate *entropy_score* during evaluation since there is no batch-specific cell population.

The divergence and entropy estimations were calculated for pairs of batches and then averaged to acquire a summary of the batch-correction performance among multiple batches. When the dimensionality of the embedding was high, *divergence_score* and *entropy_score* were calculated based on the 2dimensional UMAP [30] embeddings of the data to derive robust estimations of divergence and entropy. UMAP is a general dimensionality reduction algorithm that can achieve competitive results compared to t-SNE [50], while preserves more global structures of the data.

Third, since *divergence_score* and *entropy_score* are both proposed to evaluate the mixture of cells among batches, we also compared a metric to evaluate the separation of different cell types after batch effects being removed. To this end, we calculated the silhouette coefficient with clusters defined by cell types. For a cell **k**, let *a*(**k**) be the average distance between **k** and all the other cells within the same cluster and *b*(**k**) be the smallest average distance between **k** and all the cells in any other cluster, we define the silhouette coefficient of cell **k** as

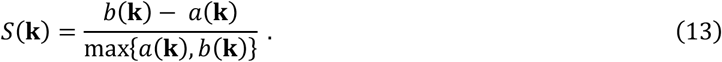

The average silhouette coefficient of all the cells from different batches is calculated after batch-effect removal, such that

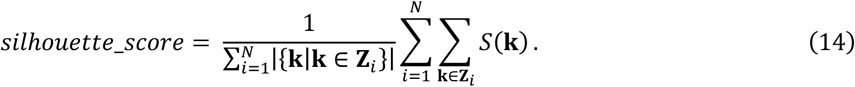

A larger *silhouette_score* indicates that the cell type assignment in the aligned dataset is more appropriate, where a cell is close to cells of the same type and distant from cells of different types. *S*(**k**) is calculated using Euclidean distance on the 2-dimensional UMAP embeddings of the results.

### Performance comparison with popular batch correction methods

We compared *BERMUDA* with several existing state-of-the-art batch correction methods for scRNA-seq data, including *mnnCorrect* [20], *BBKNN* [24], *Seurat v2* (v2.3.4) [22], *Seurat v3* (v3.0.0) [23], and *scVI*[28]. *BBKNN* and *mnnCorrect* were applied to log-transformed TPM data of variable genes. *Seurat v2, Seurat v3*, and *scVI* were applied on the datasets following the recommended workflow [22, 23, 28]. Due to the restriction of the workflow, we did not apply *Seurat v2, Seurat v3*, and *scVI* on the Gaussian simulated gene expression data. To demonstrate the necessity of batch correction methods for scRNA-seq data, we also compared *BERMUDA* with batch correction methods for microarray and bulk RNA-seq data, such as *combat* [14] and *limma* [15] (Additional file 1: Figure S3).

## List of Abbreviations

AUROC: area under the receiver operator characteristic curve
CCA: canonical correlation analysis
MMD: maximum mean discrepancy
MNN: mutual nearest neighbor
PBMC: peripheral blood mononuclear cell
RKHS: reproducing kernel Hilbert space
scRNA-seq: single-cell RNA sequencing
TMP: transcript-per-million

## Declarations

### Ethics approval and consent to participate

Not applicable.

### Consent for publication

Not applicable.

### Availability of data and materials

The implementation of *BERMUDA* can be downloaded from Github (https://github.com/txWang/BERMUDA).

### Authors’ contributions

Conceived and designed the experiments: TW, TSJ, WS. Performed the experiments and analyzed the data: TW, TSJ, ZL. Developed the structure and arguments for the paper: TW, KH. Wrote the manuscript: TW, TSJ, JZ. Edited and revised the manuscript: WS, BRH, JZ, KH. All the authors reviewed and approved of the final manuscript.

## Acknowledgements

We thank Ms. Megan Metzger for help on editing the manuscript.

## Additional files

Additional file 1: Supplementary figures and tables.

