## Supplementary information for "BERMUDA: A novel deep transfer learning method for single-cell RNA sequencing batch correction reveals hidden high-resolution cellular subtypes"

### Supplementary figures

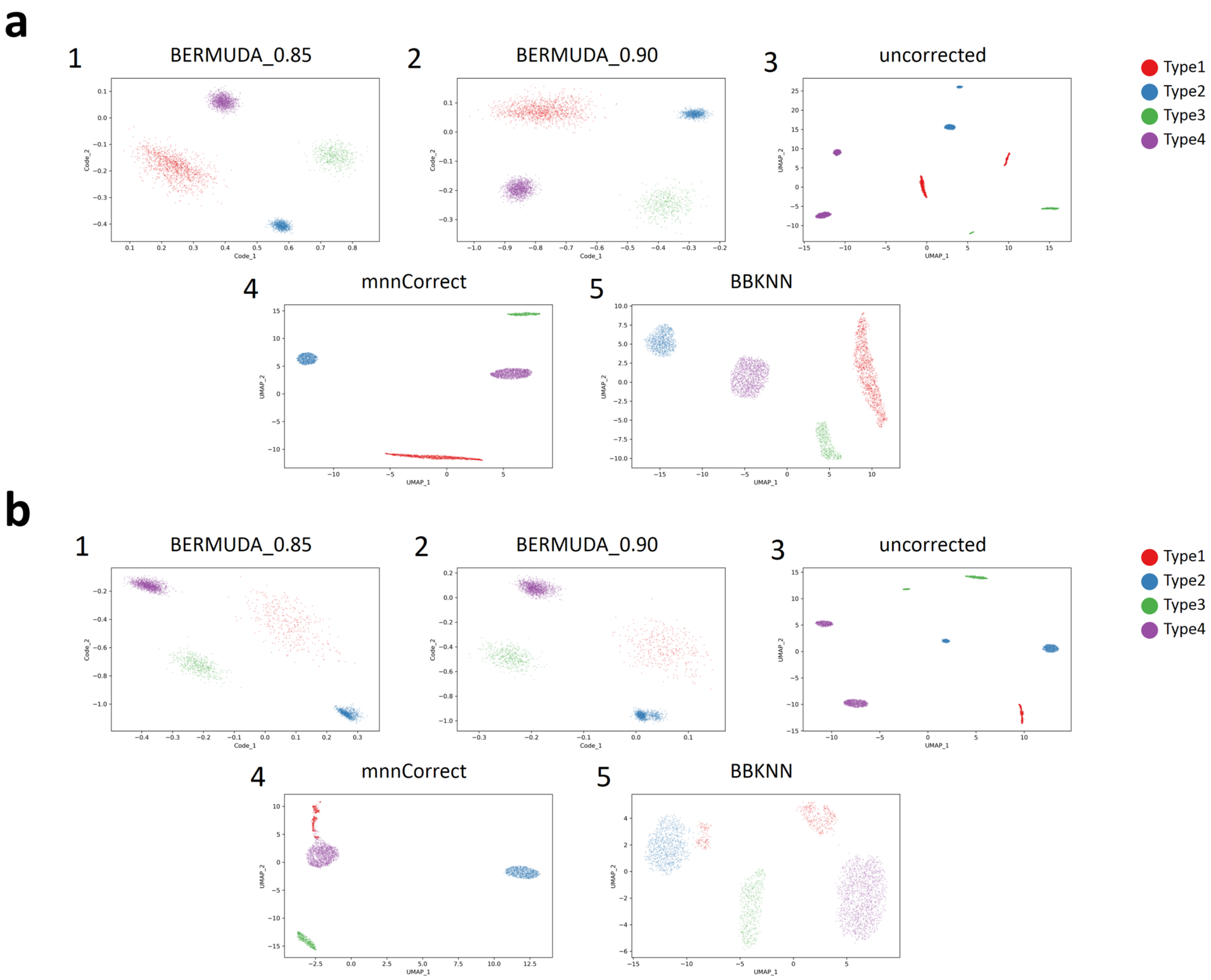


**Figure S1. Removing batch effects in simulated data generated by 2-dimensional Gaussian distribution.** Visualizations of results for simulated data generated by 2-dimensional Gaussian distribution. Results of our method are visualized by the 2-dimensional code in the trained autoencoder, while results of other methods are visualized using UMAP. BERMUDA _0.85 and BERMUDA_0.90 represent our method with $S_{thr} = 0.85$ and $0.90$ respectively. a. Results of *Experiment all*. b. Results of *Experiment removal1*.

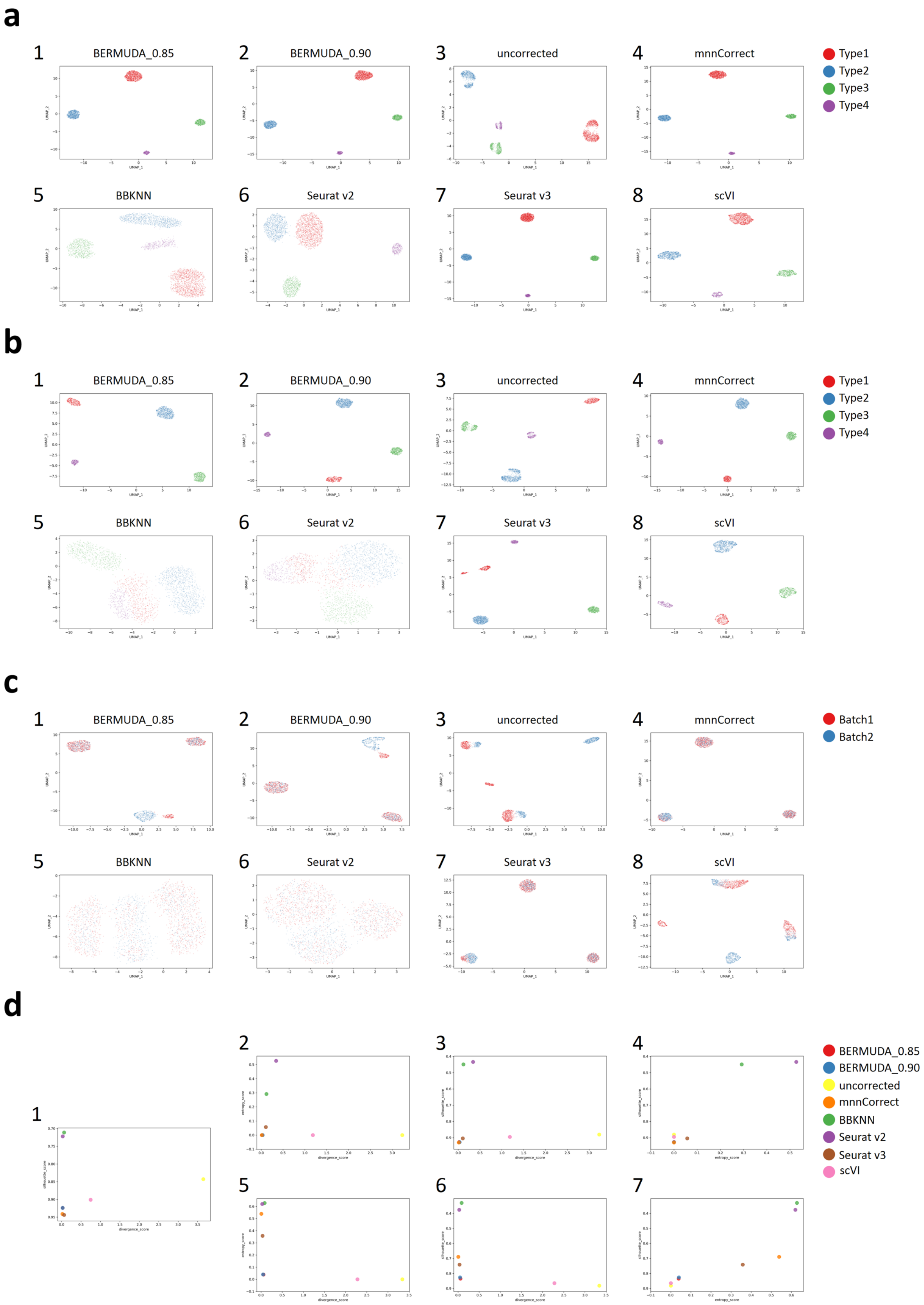
a
1 BERMUDA_0.85
2 BERMUDA_0.90
3 uncorrected
4 mnnCorrect
5 BBKNN
6 Seurat v2
7 Seurat v3
8 scVI
Type1
Type2
Type3
Type4
b
1 BERMUDA_0.85
2 BERMUDA_0.90
3 uncorrected
4 mnnCorrect
5 BBKNN
6 Seurat v2
7 Seurat v3
8 scVI
Type1
Type2
Type3
Type4
c
1 BERMUDA_0.85
2 BERMUDA_0.90
3 uncorrected
4 mnnCorrect
5 BBKNN
6 Seurat v2
7 Seurat v3
8 scVI
Batch1
Batch2
d
1
2
3
4
5
6
7
BERMUDA_0.85
BERMUDA_0.90
uncorrected
mnnCorrect
BBKNN
Seurat v2
Seurat v3
scVI

**Figure S2. Removing batch effects in simulated data generated by Splatter.** a. UMAP visualizations of results for *Experiment all* colored by cell types. b. UMAP visualizations of results for *Experiment removal1* colored by cell types. c. UMAP visualizations of results for *Experiment removal2* colored by batches. d. Evaluation of batch-correction performance on Splatter dataset using the proposed metrics. The $silhouette\_score$ axis is reversed so that points close to the bottom-left corner indicate better results. c1. *Experiment all*. c2-4. *Experiment removal1*. c5-7. *Experiment removal2*.

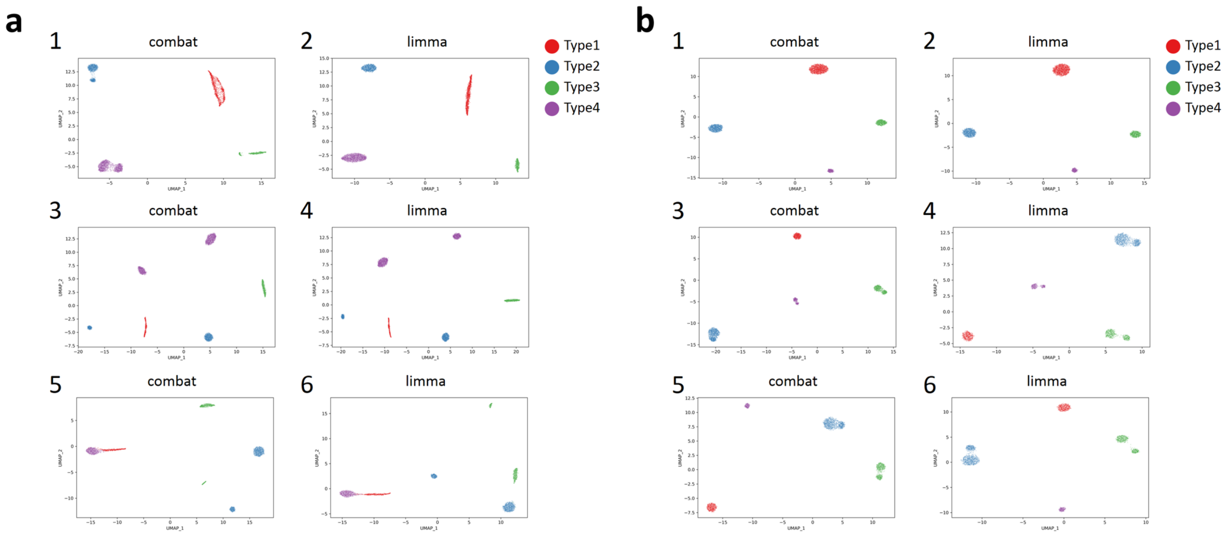


**Figure S3. *Combat* and *limma* fail to correctly remove batch effects in simulated datasets.** *Combat* and *limma* could not remove batch effects in scRNA-seq data correctly when not all cellular states were shared by all the batches (*Experiment removal1* and *Experiment removal2*). a. UMAP visualizations of batch-correction results for simulated data generated by 2-dimensional Gaussian distribution. a1-2. *Experiment all*. a3-4. *Experiment removal1*. a5-6. *Experiment removal2*. b. UMAP visualizations of batch-correction results for simulated data generated by Splatter. b1-2. *Experiment all*. b3-4. *Experiment removal1*. b5-6. *Experiment removal2*.

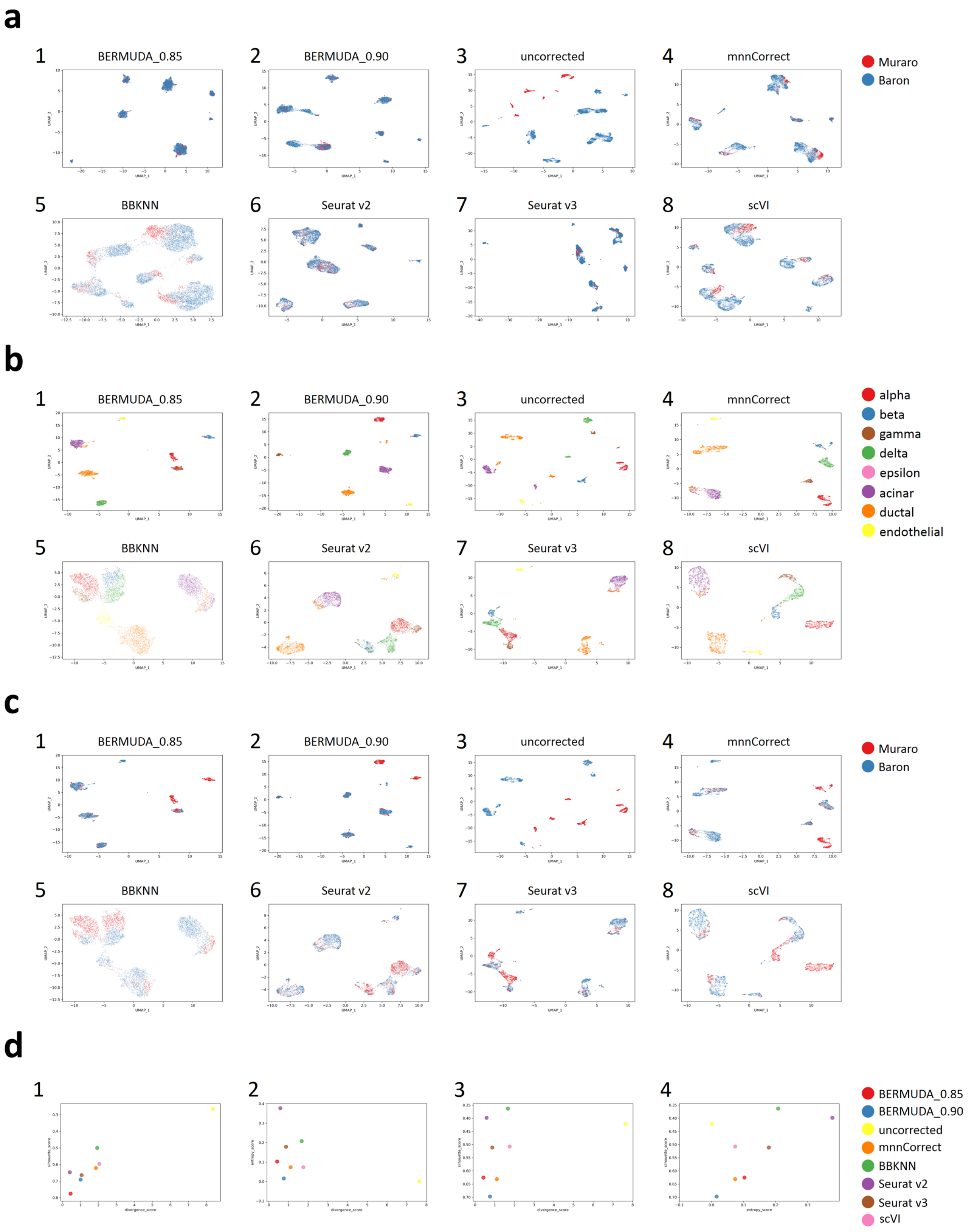


**Figure S4. Removing batch effects in scRNA-seq data of pancreas cells with *Muraro batch* and *Baron batch*.** a. UMAP visualizations of results for *Experiment all* colored by batches. b. UMAP visualizations of results for *Experiment removal* colored by cell types. c. UMAP visualizations of results in

*Experiment removal* colored by batches. d. Evaluation of batch-correction performance on *Experiment removal* using the proposed metrics. The $silhouette\_score$ axis is reversed so that points close to the bottom-left corner indicate better results. d1. *Experiment all*. d2-4. *Experiment removal*.

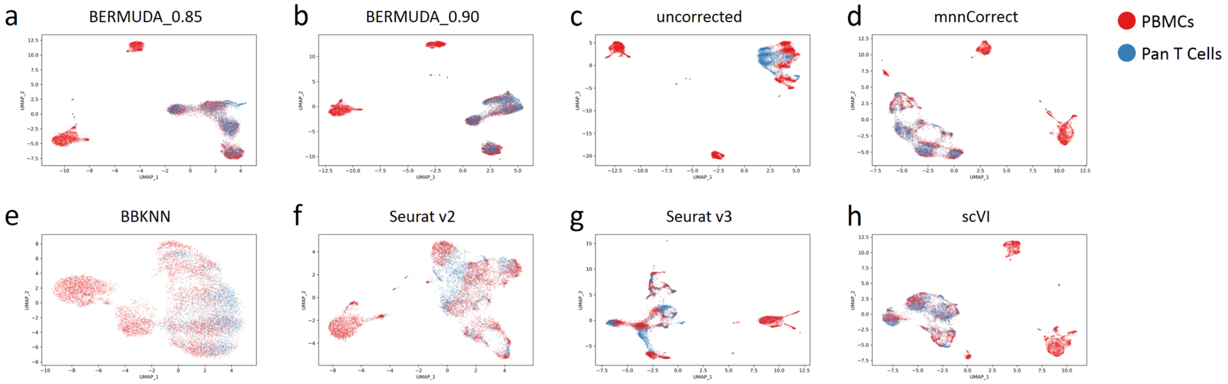


**Figure S5. Removing batch effects in scRNA-seq data of PBMCs.** UMAP visualizations of results on PBMC dataset colored by batches. Our method correctly merged T cells from the both batches, while preserved the structures of cell clusters specific to the *PBMC batch*.

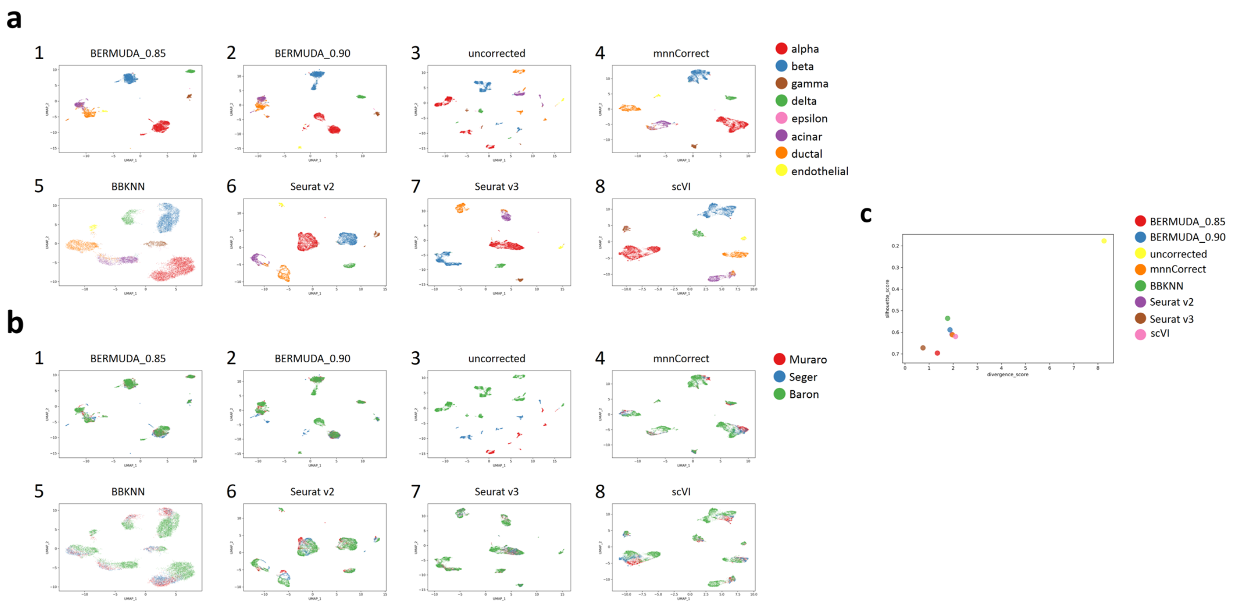


**Figure S6. Removing batch effects in scRNA-seq data of pancreas cells with multiple batches.** All the cells from *Muraro batch*, *Baron batch*, and *Segerstolpe batch* were used for analysis. a. UMAP visualizations of results for *Experiment all* colored by cell types. b. UMAP visualizations of results for *Experiment all* colored by batches. c. Evaluation of batch-correction performance on *Experiment all* using the proposed metrics.

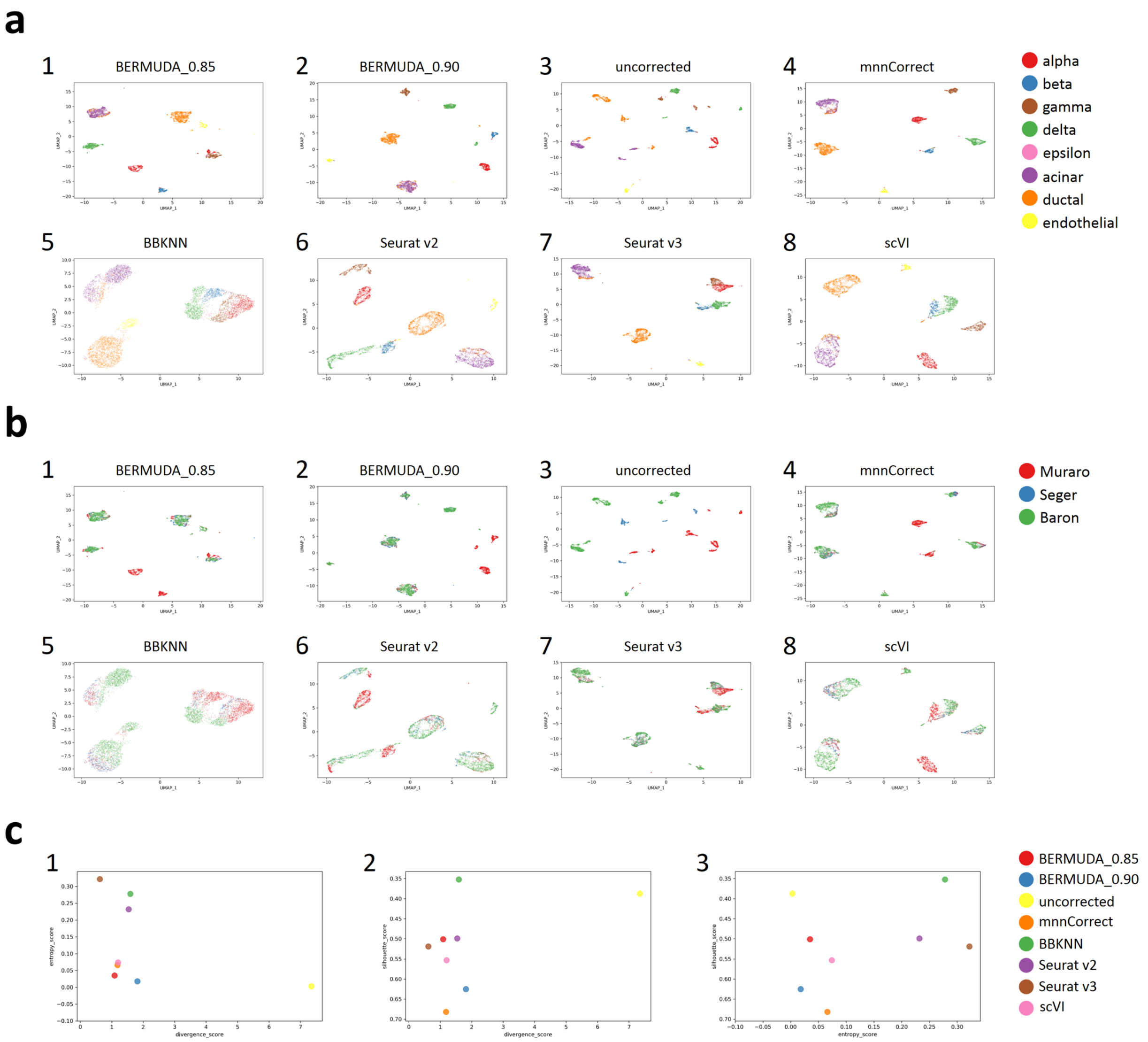


**Figure S7. Removing batch effects in scRNA-seq data of pancreas cells with multiple batches of different cell population compositions.** Alpha and beta cells from *Baron batch* and *Segerstolpe batch* were removed from analysis. a. UMAP visualizations of results for *Experiment removal* colored by cell types. b. UMAP visualization of results for *Experiment removal* colored by batches. c. Evaluation of batch-correction performance on *Experiment removal* using the proposed metrics.

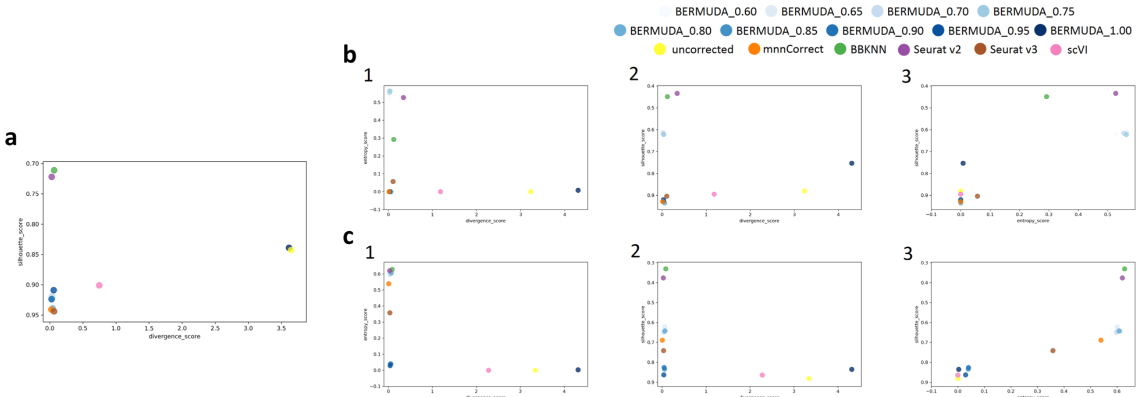


**Figure S8. Results of Splatter dataset using different $S_{thr}$ values evaluated by proposed metrics.** a. *Experiment all*. b. *Experiment removal1*. c. *Experiment removal2*. The most important parameter in our method is $S_{thr}$, which is the threshold applied on the cluster similarity score to identify similar clusters across different batches. $S_{thr}$ can affect the results of batch-correction, where a lower $S_{thr}$ value can produce a more homogeneous mixture of different batches within cell types and a higher $S_{thr}$ value can help to retain more batch-specific biological signals. We experimentally demonstrate our choice of $S_{thr}$ by evaluating the performance of our method using $S_{thr}$ ranges from 0.60 to 1.00. Our method consistently outperformed the existing methods on the Splatter dataset when choosing $S_{thr}$ between 0.85 and 0.90. More specifically, when all the cell types were shared in different batches, generally $S_{thr} \leq 0.9$ produced competitive results, since we did not need to consider the case where different cell types might be mixed together by using a low threshold. However, when we introduced large differences in cell population compositions by removing cell types from specific batches, we observed that $S_{thr}$ between 0.85 and 0.90 consistently produced best results across different experiments, and we used $S_{thr}$ between 0.85 and 0.90 as a default parameter choice for *BERMUDA*.

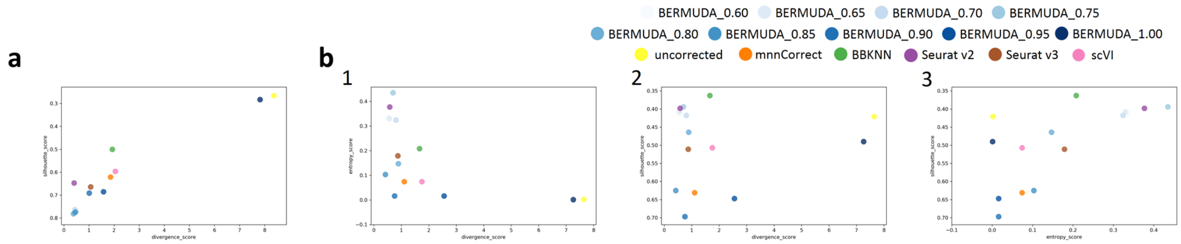


**Figure S9. Results of pancreas dataset with *Muraro batch* and *Baron batch* using different $S_{thr}$ values evaluated by proposed metrics.** a. *Experiment all*. b. *Experiment removal*. Our method consistently outperformed existing methods on the pancreas dataset when choosing $S_{thr}$ between 0.85 and 0.90.

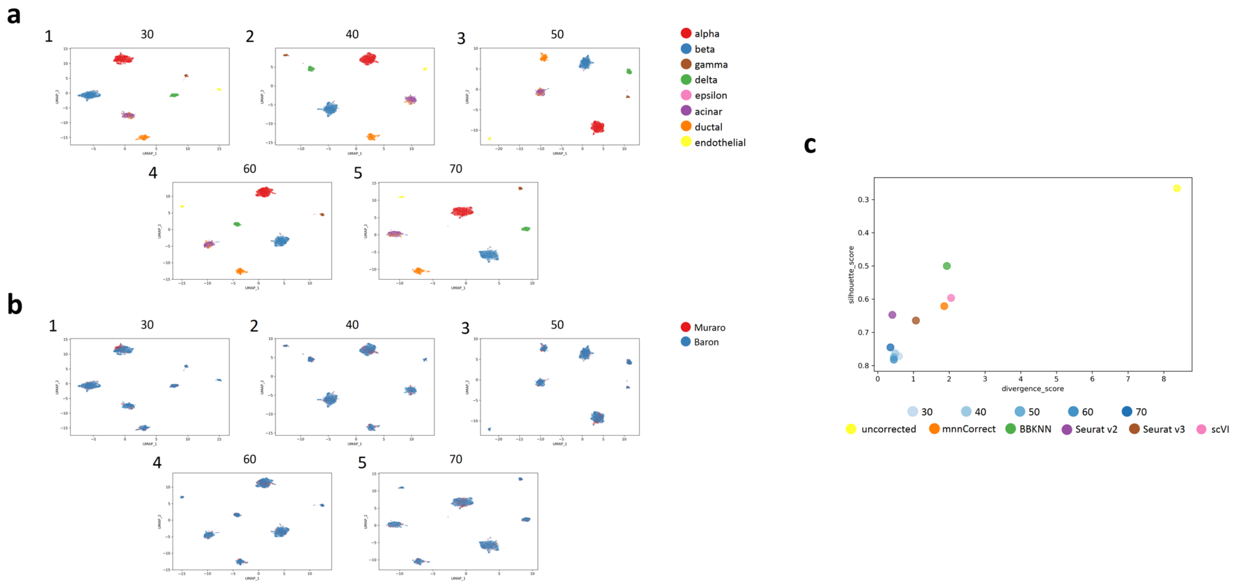


**Figure S10. *BERMUDA* is robust to the number of cells sampled from each cluster in a mini-batch.** The number of cells sampled from each cluster in a mini-batch, $n_{mb}$, is a hyperparameter in *BERMUDA*. We experimentally demonstrate that *BERMUDA* is robust to the choice of $n_{mb}$ by evaluating the performance of *BERMUDA* using $n_{mb}$ ranges from 30 to 70. The results were generated by performing *Experiment all* to combine *Muraro batch* and *Baron batch* in the pancreas dataset with $S_{thr} = 0.85$. We observed that the performance of *BERMUDA* was insensitive to the choice of $n_{mb}$ and *BERMUDA* consistently outperformed existing methods under a wide range of $n_{mb}$ values. a. UMAP visualizations of results colored by cell types. The number above each figure represents the value of $n_{mb}$ used for training *BERMUDA*. b. UMAP visualizations of results colored by batches. c. Evaluation of batch-correction performance using the proposed metrics.

### Supplementary tables

**Table S1.** List of differently expressed genes within alpha cells in the pancreas dataset.

| Gene Symbol | Adjusted p-value (Muraro vs. Baron) | Adjusted p-value (Baron vs. Baron) |
|---|---|---|
| PCSK1N | 0.00E+00 | 7.97E-55 |
| FAP | 0.00E+00 | 6.04E-76 |
| G6PC2 | 7.31E-290 | 3.54E-168 |
| SLC30A8 | 5.80E-281 | 2.75E-162 |
| SLC38A4 | 1.53E-277 | 7.67E-51 |
| ARRDC4 | 5.64E-249 | 1.88E-60 |
| ABCC8 | 6.70E-241 | 2.05E-130 |
| TM4SF4 | 1.06E-173 | 8.14E-221 |
| CRYBA2 | 1.00E-169 | 1.86E-202 |
| **MAFB** | **1.42E-158** | **4.16E-109** |
| **ARX** | **1.09E-145** | **3.21E-60** |
| TXNIP | 4.58E-104 | 6.57E-122 |
| INSIG1 | 9.34E-95 | 9.48E-53 |
| FXYD6 | 1.41E-93 | 6.77E-119 |
| XIST | 6.48E-85 | 1.19E-169 |
| SERPINA1 | 1.62E-76 | 3.09E-52 |
| S100A11 | 7.41E-74 | 7.02E-87 |
| TNFRSF12A | 1.38E-69 | 6.32E-94 |
| ANXA2 | 1.06E-64 | 1.45E-111 |

Only genes with adjusted p-value $\leq 10^{-50}$ in both tests are reported. The bold genes are discussed in detail in the paper.

**Table S2.** List of differently expressed genes within beta cells in the pancreas dataset.

| Gene Symbol | Adjusted p-value (Muraro vs. Baron) | Adjusted p-value (Baron vs. Baron) |
|---|---|---|
| SYT13 | 6.98E-194 | 2.14E-84 |
| SLC30A8 | 1.34E-179 | 2.47E-223 |
| ABCC8 | 2.05E-170 | 1.95E-162 |
| G6PC2 | 3.16E-162 | 5.89E-155 |
| INS | 3.93E-160 | 5.62E-83 |
| **PDX1** | **1.35E-151** | **8.21E-51** |
| **MAFB** | **1.06E-150** | **6.35E-95** |
| **MAFA** | **9.22E-135** | **2.77E-54** |
| PLCXD3 | 7.72E-126 | 1.97E-62 |
| PCSK1 | 1.90E-97 | 7.74E-96 |
| WNT4 | 1.11E-81 | 1.27E-61 |
| TIMP2 | 4.85E-80 | 1.90E-61 |
| EDN3 | 5.89E-62 | 4.68E-92 |

Only genes with adjusted p-value $\leq 10^{-50}$ in both tests are reported. The bold genes are discussed in detail in the paper.

**Table S3.** Running time of different methods in the pancreas dataset.

| Method | Running time(s) |
|---|---|
| BERMUDA ($S_{thr} = 0.85$) | 285.91 |
| BERMUDA ($S_{thr} = 0.90$) | 262.98 |
| mnnCorrect | 35.90 |
| BBKNN | 5.02 |
| Seurat v2 | 338.33 |
| Seurat v3 | 75.27 |
| scVI | 360.33 |

The running time was measured by performing *Experiment all* on *Muraro batch* and *Baron batch*. Since we expect *BERMUDA* to be adopted by biologists who may not always have easy access to high-end computing facilities, and some of the methods compared do not have a GPU implementation, we evaluated the running time on a desktop computer with a CPU (2.7 GHz Intel Core i5) for fair comparison.
